# Simultaneous mapping of epigenetic inter-haplotype, inter-cell and inter-individual variation via the discovery of jointly regulated CpGs in pooled sequencing data

**DOI:** 10.1101/2023.02.28.530419

**Authors:** Benjamin Planterose Jiménez, Brontë Kolar, Manfred Kayser, Athina Vidaki

**Affiliations:** Department of Genetic Identification, Erasmus MC, University Medical Center Rotterdam, Rotterdam, the Netherlands

**Author notes:** Correspondence: Benjamin Planterose Jiménez, Primary, Secondary.

## Abstract

In the post-GWAS era, great interest has arisen in the mapping of epigenetic inter-individual variation towards investigating the emergence of phenotype in health and disease. Relevant DNA methylation methodologies – epigenome-wide association studies (EWAS), methylation quantitative trait *loci* (mQTL) mapping and allele-specific methylation (ASM) analysis – can each map certain sources of epigenetic variation and all depend on matching phenotypic/genotypic data. Here, to avoid these requirements, we developed Binokulars, a novel randomization test that identifies signatures of joint CpG regulation from reads spanning multiple CpGs. We tested and benchmarked our novel approach against EWAS and ASM on pooled whole-genome bisulfite sequencing (WGBS) data from whole blood, sperm and combined. As a result, Binokulars simultaneously discovered regions associated with imprinting, cell type- and tissue-specific regulation, mQTL, ageing and other (still unknown) epigenetic processes. To verify examples of mQTL and polymorphic imprinting, we developed JRC_sorter, another novel tool that classifies regions based on epigenotype models, which we deployed on non-pooled WGBS data from cord blood. In the future, this approach can be applied on larger pools to simultaneously map and characterise inter-haplotype, inter-cell and inter-individual variation in DNA methylation in a cost-effective fashion, a relevant pursuit towards phenome-mapping in the post-GWAS era.

## Introduction

Genome-wide association studies (GWAS) have enabled the discovery of a myriad of genetic variants influencing human traits in health and disease. However, almost two decades later, most of the human phenome remains largely unexplained by solely genetic variants. Moreover, discovered genetic associations, which frequently map to the non-coding genome, remain to be rationalized by mechanistic models (Do et al. 2017; Brandes et al. 2022). One level higher, the epigenome lies in the interphase between the genome and the environment. Thus, its study may help elucidate relevant environmental factors influencing human traits that remain unexplained by genetic variants. Additionally, it may unveil mechanisms of action for genetic associations that mediate their effect via the epigenome. In particular, DNA methylation is the most characterized epigenetic biomarker and plays a key role in development, genome regulation and organization. For instance, it is involved in genomic imprinting, X-inactivation, the suppression of selfish elements and ageing (Mattei et al. 2022). In mammals, DNA methylation almost exclusively occurs in the form of 5-methylcytosine at the context of CpG sites (cytosine followed by guanine in 5’→3’ orientation) (Sarkies 2022). Driven by all of the above, there is currently a growing interest in mapping inter-individual variation of human CpG methylation to different genetic and environmental factors (Garg et al. 2018; Gunasekara et al. 2019).

Towards this aim, several approaches have been developed: i) epigenome-wide association studies (EWAS), ii) methylation quantitative trait *loci* (mQTL) mapping and iii) allele-specific methylation (ASM) analysis (Flanagan 2015; Villicaña and Bell 2021). In EWAS and mQTL mapping, the methylation of individual or groups of CpGs are tested for association with a particular identifiable phenotypic trait or the genotype of particular genetic variants, respectively (Michels et al. 2013; Villicaña and Bell 2021).

To achieve highly powered studies, both EWAS and mQTL mapping mostly quantify DNA methylation via microarrays (Michels et al. 2013). However, this comes at the cost of ignoring most of the methylome: the two popular platforms – Infinium HumanMethylation450 and MethylationEPIC Beadchip microarrays – target only 1.5 % and 3 % of the methylome, respectively. Additionally, one has to consider various technical issues such as background fluorescence (Triche et al. 2013), batch effects (Ross et al. 2022), genetic artifacts (Planterose Jiménez et al. 2021) or cross-hybridizing probes (Chen et al. 2013; Price et al. 2013). Only recently, EWAS and mQTL mapping have been carried out on whole-genome bisulfite sequencing (WGBS) data – the gold standard of methylomics – but whose popularity has remained largely quenched due to its high cost and DNA input requirements (Suzuki et al. 2018; Mordaunt et al. 2020; Perzel Mandell et al. 2021). In contrast to EWAS and mQTL mapping, ASM analysis is mostly performed on a single WGBS experiment by relying on heterozygous genetic variants; howbeit, microarray implementations for this assay used to be popular in the past (Kerkel et al. 2008). In its modern version, ASM tests for allelic asymmetries by jointly measuring SNP and CpG (epi)alleles across reads that include both markers. While ASM has been traditionally used to discover imprinted *loci* (whose methylation is associated with parent-of-origin) by using SNPs to label each haplotype, it can also discover “local” mQTL (Tycko 2010): when in close proximity, a bi-allelic *cis*-acting causal variant (or a variant in linkage disequilibrium (LD) with it) can be co-sequenced in the same read alongside with CpGs belonging to the co-methylation window of the mQTL.

Despite being less common in the literature, sample pooling approaches can be used as supplement to improve effective sample size, without dramatically increasing associated sequencing costs and whilst preserving the privacy of individuals in the pool. In pooled EWAS (pEWAS), cases or controls samples are pooled prior to WGBS (Docherty et al. 2010). In pooled ASM (pASM), local mQTL associations are enriched by depleting spurious parent-of-origin/allele associations (Kaplow et al. 2015). In perspective, all discussed approaches (EWAS/pEWAS, mQTL mapping, ASM/pASM) require profiling additional information (genotype, phenotype) prior to mapping inter-individual variation in the methylome. Here, we argue that this step is not strictly necessary to identify signatures of active DNA methylation regulation. Though the human genome contains over 30 million CpG sites, these are not regulated independently due the natural processivity of the enzymes involved in DNA methylation regulation, namely DNA-methyltransferases (DNMT) 1, 3A, 3B and ten-eleven translocation (TET) enzymes (Vilkaitis et al. 2005; Holz-Schietinger and Reich 2010; Fu et al. 2012; Xing et al. 2020); this is the ability to perform several catalytic cycles without detaching from DNA. In other words, the CpG level is not the ultimate unit of DNA methylation regulation resolution in the human genome (Guo et al. 2017).

In this study, we re-examine current approaches to map inter-individual variation in the human methylome. In particular, we highlight the ability of a pooled WGBS experiment to simultaneously capture inter-haplotype, inter-cell and inter-individual variation in a cost-effective and privacy-preserving fashion. Importantly, WGBS data provides an opportunity to explore the joint probability distribution between the epialleles of neighbouring CpGs within the same read. We define co-methylation within reads (CWR) as the tendency for CpGs within the same read to display the same epiallele (unmethylated/methylated) at a higher level than what it is expected by chance. Using a coin toss as an analogy to the null hypothesis model (head: methylated; tails: unmethylated), a read with *n* CpGs corresponds to *n* coin tosses where the number of heads (i.e. number of methylated cytosines) follows a binomial distribution with parameter *p* (i.e. methylation ratio of a given region). However, in principle, active DNA methylation regulation breaks the assumption of independence between neighbouring CpGs: processive runs of DNMT or TET favours the emergence of reads where all CpGs in a row are fully methylated or unmethylated, respectively. Thus, in theory, we can identify active regulation by identifying regions which concentrate reads that deviate from this null hypothesis. This would enable the simultaneous mapping of DNA methylation inter-haplotype, inter-cell or inter-individual variation and thus, integrating a wide range of epigenetic processes.

Given this promising prospect, the aim of our study was to identify jointly regulated CpGs (JRCs) using pooled WGBS data. Towards this end, we developed Binokulars (**Bino**mial li**k**elihood f**u**nction-based bootstrap hypothesis test for co-methy**la**tion within **r**ead**s**), a novel statistical test for detecting CWR. As part of our simulation benchmark, we confirmed its favourable performance even at low coverage, alongside considerable improvements with increasing CpG density. For effective genome-wide testing, we also developed JRC_seeker, a novel pipeline wrapper that incorporates hidden Markov models (HMMs) to pre-select intermediately methylated regions (IMRs) to test with Binokulars. We deployed JRC_seeker on pooled WGBS data in whole blood (WB), sperm (SP) and the combination of both and contrasted our results with those obtained with pEWAS and pASM. Following this strategy, we simultaneously discovered JRCs associated with a wide range of phenomena: imprinting, cell type- and tissue-specific regulation, mQTL and ageing. Finally, we developed another novel tool, JRC_sorter, that can classify JRCs into five epigenotype models given non-pooled WGBS data. With this tool and using data in cord blood, we additionally verified JRCs discovered in WB, as examples of mQTL and polymorphic imprinting.

## Results

### Identification of co-methylation within reads with Binokulars

We developed Binokulars (**Bino**mial li**k**elihood f**u**nction-based bootstrap hypothesis test for co-methy**la**tion within read**s**), a randomization test for CWR (co-methylation within reads) from pooled WGBS data. Binokulars operates at the level of reads to identify signatures of CpG joint regulation (**Fig. 1A-B**, more details in **Supplementary Methods: Section I**). We hereon refer to significant regions identified by Binokulars as JRCs (jointly regulated CpGs). To understand its limits, we firstly evaluated Binokular’s performance via a simulation study (**Fig. 1C**). The scope and limitations of our simulations are discussed at **Supplementary Methods: Section II.** This way, we verified that Binokulars can identify JRCs with great power, especially at a high CpG density (when it is more likely to obtain reads with several CpGs). Moreover, Binokulars has very low allelic frequency and coverage requirements, making it optimal for deployment on sparse WGBS data (**Fig. 1C**).

**Figure 1.**
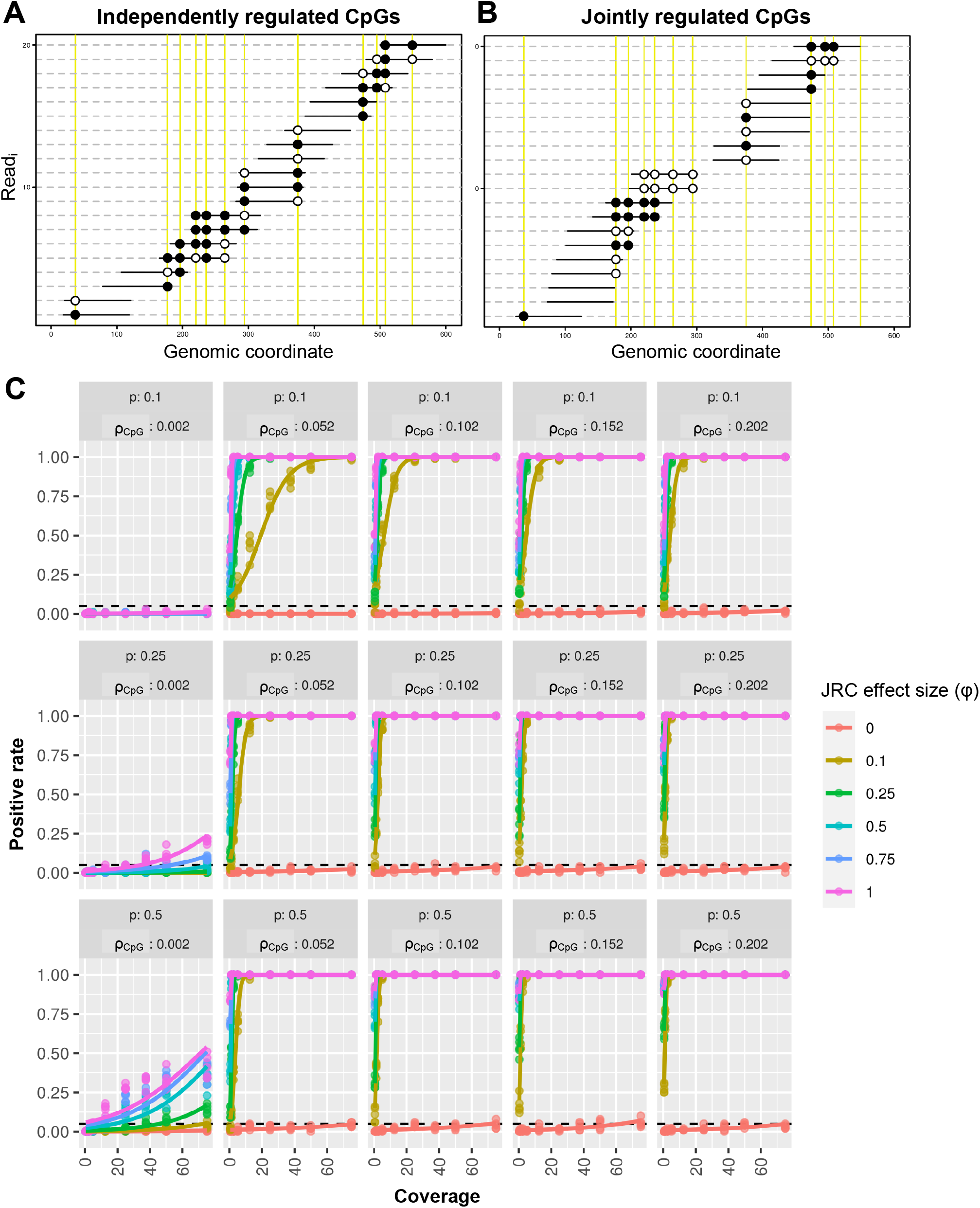
Simulating JRCs to test Binokulars. Simulated example of a CpG with (A) non-JRC (effect size (φ) = 0) and (B) JRC behaviour (φ = 1). CpG placement positions are coloured in yellow. (B) Performance of Binokulars as a function of coverage, epiallele frequency (p), CpG density (ρ_CpG_) and φ at fixed region length (L) = 2000 and read length (l) = 100. Binokulars was tested with parameters bin size (δ)=200 and number of bootstraps (N_boot) = 1000. We employed positive rate (PR) as the performance metric; equal to the true positive rate (TPR) when φ ≠ 0 or equal to false positive rate (FPR) when φ = 0. To compute PR, a total of 100 iterations were performed, each with a different CpG placement. To assess PR’s error, we additionally computed 5 replicates per condition. The fitted lines correspond to a generalized linear models (glm) with a quasi-binomial link function.

### Epigenome-wide identification and functional annotation of jointly regulated CpGs

Binokulars tests whether an input region behaves as a JRC given pooled WGBS data. However, running such a randomization test at the level of reads cannot be trivially scaled genome-wide due to its high computational burden. Yet, significant relief can be achieved by preselecting IMRs (intermediately methylated regions) across the WGBS pool, as fully methylated/unmethylated regions comprise a substantial subset of the genome that is of no use for JRC discovery. This task is fit for HMMs (hidden Markov models). We employed the implementation by ChromHMM (Ernst and Kellis 2017). Additionally, to solve the conflict between the number of required permutations and the genome-wide multiple testing burden, we proposed a parametric approximation for the permuted test statistic under the null distribution. The combination of these elements gave rise to JRC_seeker, a Snakemake pipeline (Köster and Rahmann 2012) that enables genome-wide deployment of Binokulars (**Fig. S2**).

As a proof-of-principle, we applied JRC_seeker on WGBS data from pooled WB (whole blood), SP (sperm) and pooled whole blood+sperm (WB+SP) derived from 12 healthy males. Our choice of tissues and samples is pertinent. Firstly, WB is a highly heterogeneous tissue as oppose to SP; thus, the WB pool includes cell-type methylation variation. Secondly, the SP pool is composed by solely male haplotypes and thus, does not include parent-of-origin methylation variation (imprinting). Thirdly, the WB+SP pool is additionally subject to tissue-specific methylation regulation. Fourthly, since all samples were derived from males, neither sex nor X-inactivation methylation variation is present in any of the pools. Finally, the inclusion of individuals from a broad age range (50 % aged 18-24 and 50 % aged 61-71) means that ageing variation is present in all pools. In the case of WB, this also includes cell-type composition that changes with age (Bergstedt et al. 2022). We performed quality control at four different levels: i) sequencing reads (**Fig. S3**), ii) bisulfite alignment (**Fig. S4**), iii) genome segmentation (**Fig. S5**) and iv) segmentation polishing (**Fig. S6-8**). Corresponding Manhattan plots, QQ-plots and p-value distributions are available (**Fig. S9-11**).

We discovered a total of 4215 (WB), 7386 (SP) and 56,896 (WB+SP) JRCs from a total of 662,659 (WB), 536,691 (SP) and 667,071 (WB+SP) IMRs at Bonferroni significance threshold, with corresponding chromosomic and size distributions (**Fig. S12**). From here onwards, we refer to these significant sets as JRC^WB^, JRC^SP^, JRC^WB+SP^, respectively. To highlight a few examples, the top JRC^WB^ corresponds to a region at intron 2 from the imprinted gene retinoblastoma 1 (*RB1*) (**Fig. 2A**), previously highlighted for its relevance in oncology (Buiting et al. 2010). In fact, imprinted genes were strongly overrepresented among JRCs^WB^ (p-val = 7.1×10^-16^, Fisher’s exact test) and slightly enriched at JRCs^WB+SP^(p-val = 0.003, Fisher’s exact test), as supposed to JRCs^SP^(p-val = 0.43, Fisher’s exact test). JRCs^WB^were associated with 39 imprinted genes, including pivotal examples like insulin like growth factor 2 (*IGF2)*, guanine nucleotide binding protein, alpha stimulating (*GNAS)*, Lethal 3 malignant brain tumor-like protein (*L3MBTL1)*, nucleosome assembly protein 1 like 5 (*NAP1L5)*, distinct subgroup of the Ras family member 3 (*DIRAS3)* and maternally expressed 3 *(MEG3)*. Furthermore, we found JRCs^WB^ associated with cell types: for example, at an intron of the forkhead box k1 (*FOXK1)* gene, hypo- and hyper-methylated in the myeloid and lymphoid lineage, respectively (**Fig. 2B**). We also identified JRCs^WB^associated with inter-haplotype variation, such as the intronic mQTL at the ninjurin 2 (*NINJ2*) gene (**Fig. 3A, S13**) (Planterose Jiménez et al. 2021).

**Figure 2.**
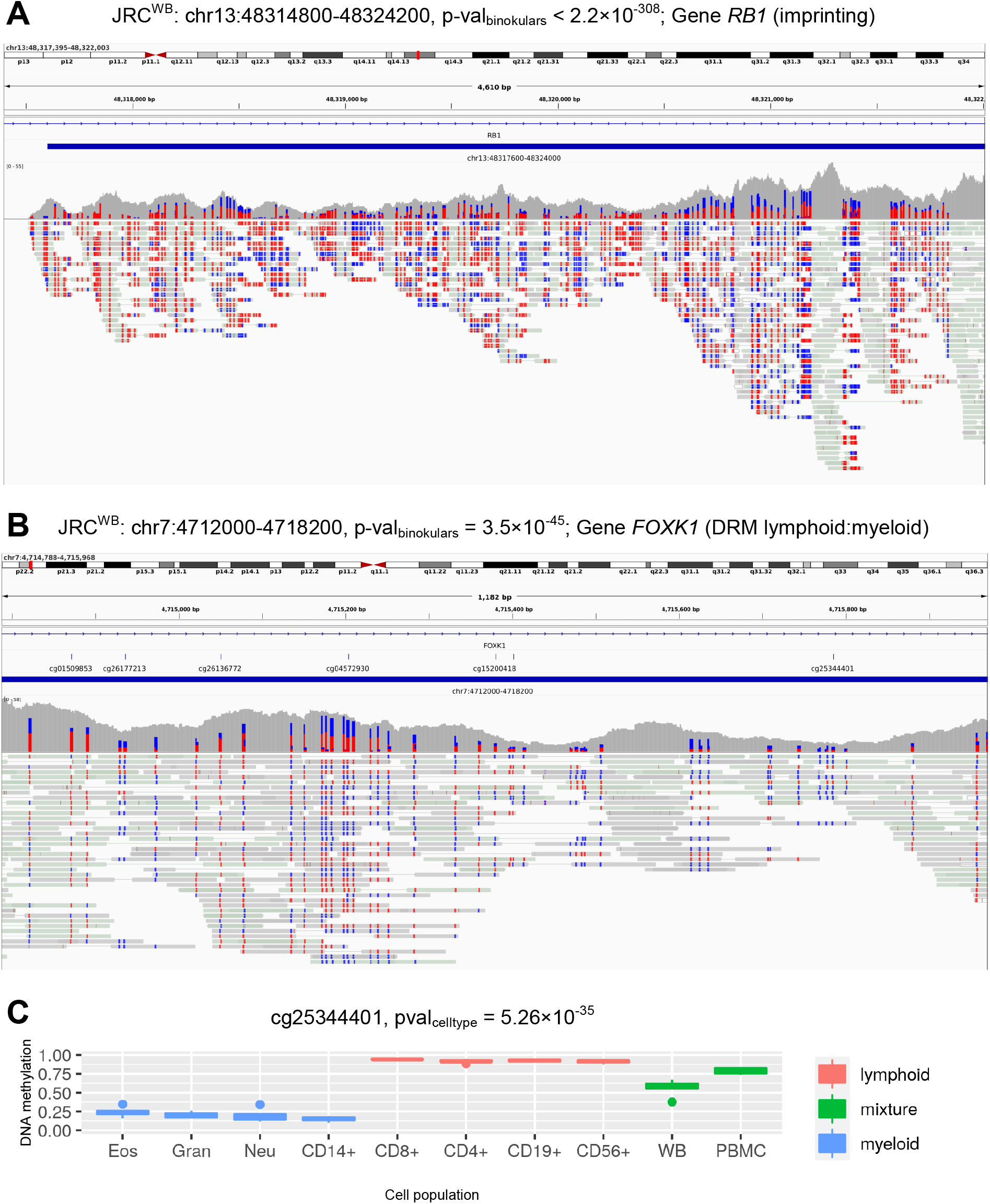
Highlighted examples of JRCs^WB^. (A) Genomic screenshot of a JRC^WB^ at an imprinted intron in the *RB1* gene. (B) Genomic screenshot of a lymphoid/myeloid lineage-associated JRC^WB^ at an intron in the *FOXK1* gene. Blue and red colouring at the level of reads represent unmethylated and methylated cytosines, respectively. (C) Cell-type composition summary statistics and methylation distributions for the probe cg25344401, in representation for the *FOXK1* JRC highlighted on panel (B), based on microarray data from FACS-sorted blood cell types. Eos: eosinophile; Gran: granulocytes; Neu: neutrophil, CD14+: monocytes, CD8+: cytotoxic T-cell; CD4+: helper T-cell; CD19+: B-cell; CD56+: NK cell; WB: Whole blood; PBMC: peripheral blood mononuclear cells.

**Figure 3.**
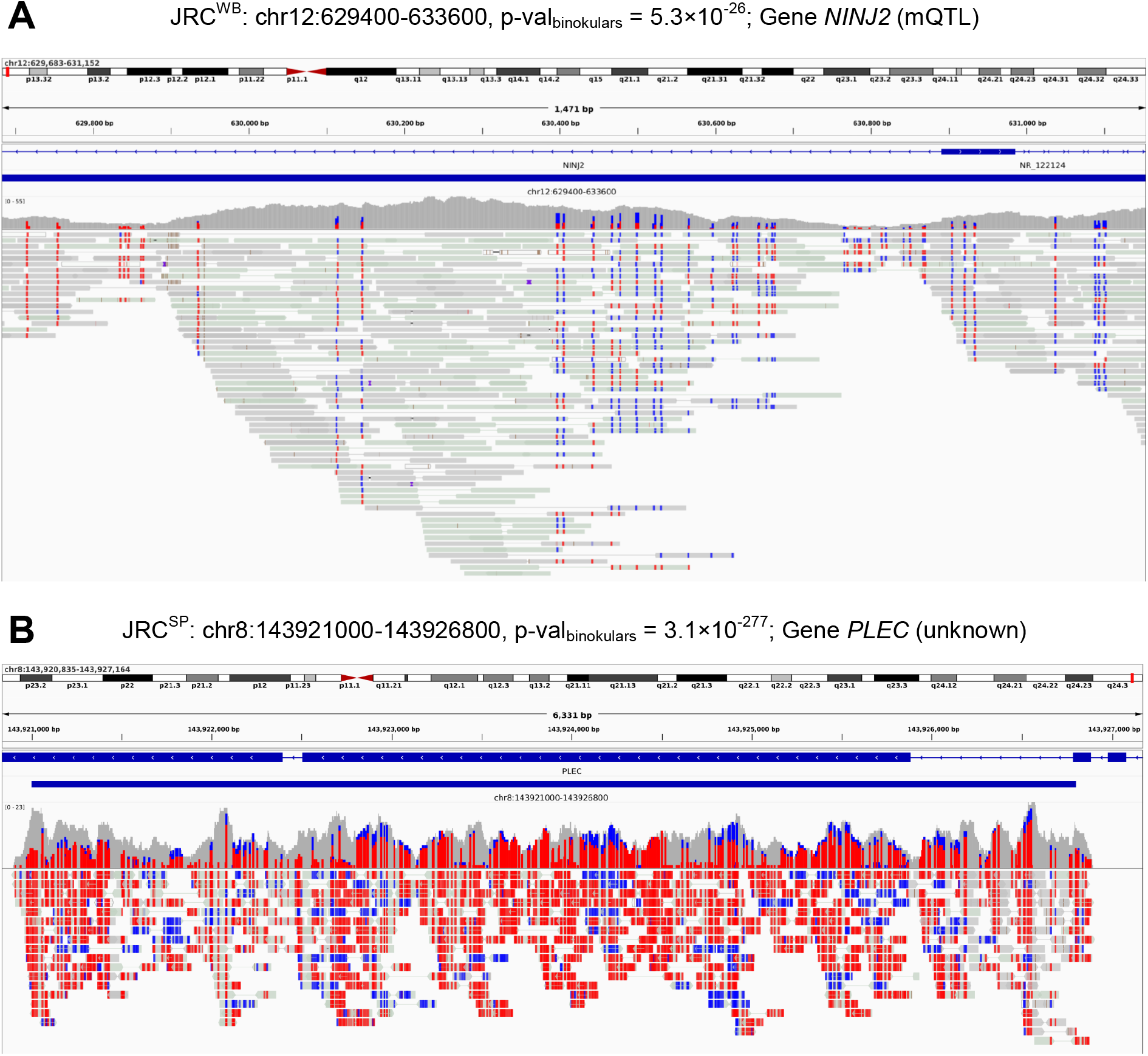
Genomic screenshots for highlighted JRC examples. (A) JRC^WB^ from a previously verified mQTL at an intron in the *NINJ2* gene. (B) JRC^SP^ at the 5’-end of the *PLEC* gene of unknown functional origin. Blue and red colouring at the level of reads represent unmethylated and methylated cytosines, respectively.

In the case of JRCs^SP^ examples, we found strong evidence for CWR covering the last two exons of the plectin (*PLEC*) gene, which we could verify across platforms but not attribute to any known source of methylation variation (**Fig. 3B, S14**). Additionally, though there has not been any mQTL mapping study in sperm as of yet to the best of our knowledge, we managed to verify the archetypical signatures of mQTL (e.g. trimodal methylation distribution) for several JRC^SP^ on DNA methylation microarray data in SP (**Fig. S15**). As for JRCs^WB+SP^, we highlight an example at the first intron of the disco-Interacting Protein 2 Homolog C (*DIP2C*) gene, which displays strong tissue-specific regulation (**Fig. 4A**).

**Figure 4.**
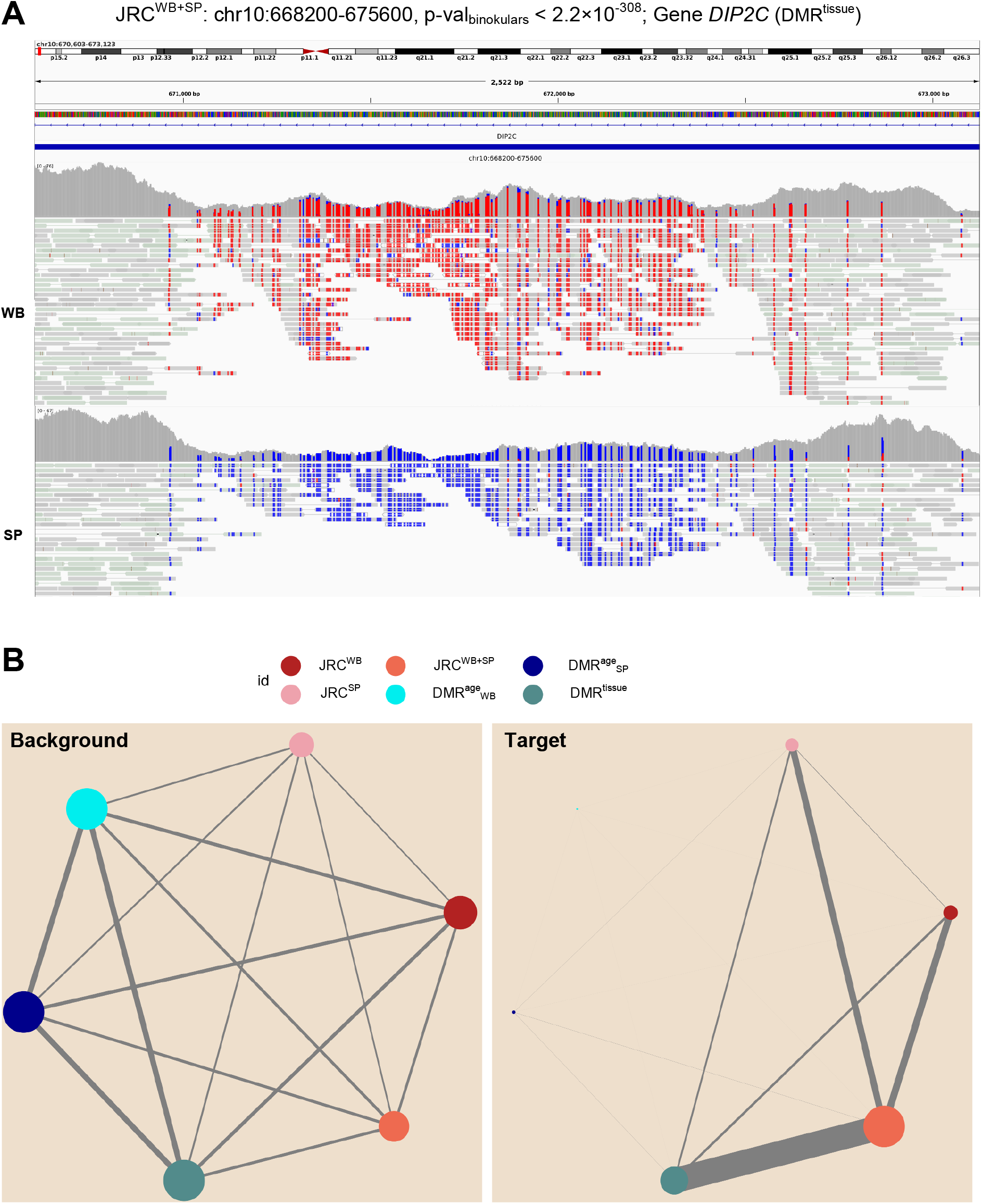
Highlighted example of JRC^WB+SP^. (A) Genomic screenshots of a JRC^WB+SP^ at an intron in the *DIP2C* gene, displaying strong inter-tissue methylation differences between WB and SP. Blue and red colouring at the level of reads represent unmethylated and methylated cytosines, respectively. (B) Colocalization via network representation of genomic sets obtained via binokulars: WB (JRC^WB^), SP (JRC^SP^) and WB+SP (JRC^WB+SP^); or pEWAS: age in WB (DMR^age^ _WB_), age in SP (DMR^age^_SP_) and WB Vs SP (DMR^tissue^). Node size is proportional to the number of megabases spanned by a given set, whilst edge size is proportional to the number of megabases shared between two genomic sets. The background set corresponds to the set of genomic regions prior to discovery, whilst the target corresponds to solely significant regions.

As part of our functional annotation, we linked JRCs to different gene features (promoter, intron, exon and intergenic) and tested for enrichment. JRCs^WB^ and JRCs^WB+SP^ were significantly enriched in promoter and exon features, whilst depleted in intron and inter-genic features (**Fig. S16-17**). Elsewise, JRCs^SP^ were significantly enriched in promoter, intergenic and exon features and depleted in intron features (**Fig. S18**). We also performed functional ontology enrichment analysis on those JRCs that could be assigned to a gene (∼ 50% for each set were non-intergenic JRCs). As a result, JRCs^WB^ and JRCs^WB+SP^were strongly enriched for a wide range of transcription factor motifs and (**Fig. S19-20**), whilst JRCs^SP^ were enriched for neurodevelopmental and neuronal GO terms (**Fig. S21**). The latter probably results from biases in GO term enrichment arising from incomplete SP ontologies, since this tissue remains critically understudied in the literature (Gaudet and Dessimoz 2017; Åsenius et al. 2020).

### Comparison of Binokulars’ findings with pooled epigenome-wide association studies

To better understand the sources of epigenetic variation that drive JRCs and for the sake of method comparison and benchmark, we performed a series of pooled EWAS on the same dataset used for discovery. To do so, we made use of the routines from the DSS R-package that allow for DMR (differentially methylated region) calling without technical replicates by integrating chromosomic spatial information. We performed three comparisons: i) EWAS for tissue; WB *versus* SP (DMR^tissue^) and EWAS for age; younger *versus* older donors in ii) WB (DMR^age^_WB_) and iii) SP (DMR^age^_SP_). For the sake of clarity, comparison i) may reveal *loci* under tissue-specific regulation or imprinting, while comparison ii) and iii) may reveal *loci* associated with age in WB (both changes in cell type composition with age and intrinsic changes with age) and SP (solely intrinsic changes with age), respectively. In total, we found 118,864 (DMR^tissue^), 135 (DMR_WB_^age^) and 1826 (DMR_SP_^age^) DMRs significantly associated for each comparison (**Fig. S22**). Examples of each set are highlighted in **Fig. S23-25**. Comparing these sets with those obtained by Binokulars should be done in caution, since their background sets are very different. Hence, we used a network visualization in which the node sizes represent the number of Mbs spanned by each set, whilst the edges represent the shared Mbs between pairs of sets (**Fig. 4**). Most notably, we found strong co-localization between DMR^tissue^ and JRC^WB+SP^, which is expected since both include regions subject to between-tissue and imprinting variation. Neatly, we also found strong co-localization between JRC^WB^ and JRC^WB+SP^ as well as between JRC^SP^ and JRC^WB+SP^. This indicates that tissue-specific JRCs can be identified in spite of having pooled different tissues altogether; though, it can also be partly explained by the co-localization between DMR^tissue^ and JRC^SP^ or JRC^WB^. To highlight some examples, we verified JRCs that were also pEWAS hits for age. We discovered that the promoter of homeobox A4 (*HOXA4*) and the lamin B2 (*LMNB2*) genes were DMRs in WB and SP, respectively (**Fig. 5**). These findings – association with age, effect size and directionality – were additionally verified on DNA methylation microarray data for WB and SP, respectively (**Fig. S26-27**).

**Figure 5.**
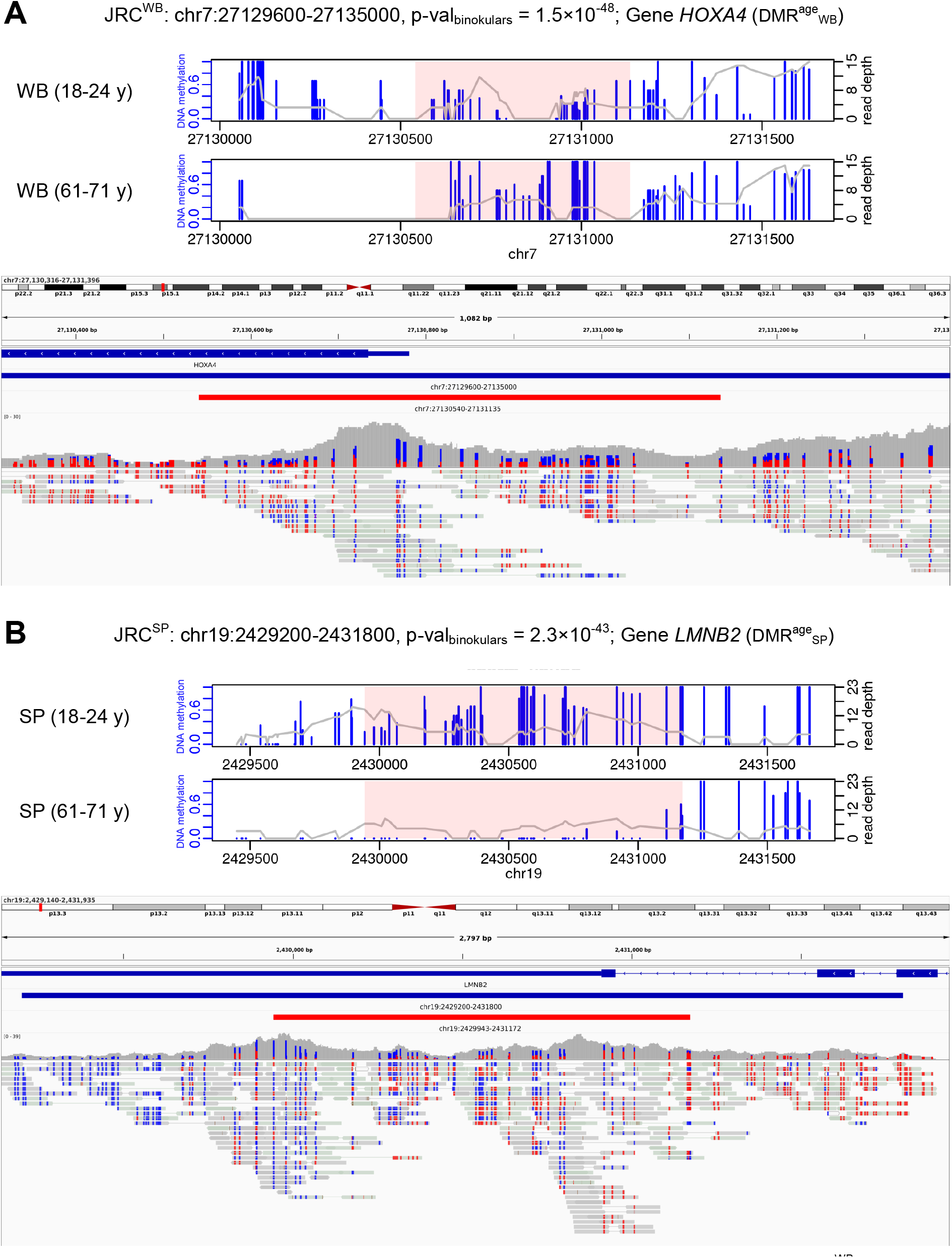
Genomic screenshots for highlighted age-associated JRC examples. (A) Age-associated JRC^WB^ at the pro-moter of the *HOXA4* gene. (B) Age-associated JRC^SP^ at the *LMNB2* gene. Blue and red colouring at the level of reads represent unmethylated and methylated cytosines, respectively.

### Comparison of Binokulars’ findings with pooled allele-specific methylation

Additionally, to systematically identify JRCs whose methylation variation can be explained by genetic variants, we carried out a pooled ASM analysis by using BISCUIT (Zhou 2022) on the same discovery dataset. In total, we tested a total of 53,709,032 (WB) and 38,299,379 (SP) heterozygous SNP-CpG pairs, respectively. However, our analysis revealed that the vast majority of comparisons were somehow compromised. To begin with, only 8.6 % (WB) and 9.5 % (SP) of the called SNPs were registered variants in dbSNP155 (MAF > 0.05), respectively. In its defence, *de novo* calling heterozygous genetic variants from low-coverage WGBS data supposes a big computational challenge (Liu et al. 2012). On top, our data was particularly demanding and error-prone due to its pooled nature. In any case, Biscuit’s sub-optimal precision in variant calling has been reported elsewhere (Lindner et al. 2022). Moreover, most of the strongly associated CpG/SNP pairs could be explained by several types of genetic artifacts caused by CpG-SNPs and indel-SNPs (more details at **Supplementary Methods: Section III, Fig. S28**). For example, a G>H SNP at position 2 of a CpG site, where H: A/C/T, causes a CpG/CpH transition (type-IIA genetic artifact). CpH methylation is almost inexistent for the vast majority of tissues in the context of mammalian cells; thus, artifactual ASM signatures arise from type-IIA artifacts when located at hyper-methylated *loci*. Though prior studies claim that these are of functional relevance (Shoemaker et al. 2010), we denote them as artifacts: CpG/CpH transitions do not necessarily induce methylation changes that propagate through the genomic neighbourhood; or at the least, it cannot be proven by the association of a single CpG/SNP pair.

Filtering out our categories of genetic artifacts resulted in solely 68 (WB) and 56 (SP) ASM pairs meeting Bonferroni significance threshold (applied post-filtering); or a total of 62 (WB) and 54 (SP) unique CpGs and 61 (WB) and 53 (SP) unique SNPs, respectively. Surprisingly, there were 21 common CpGs between WB (32.2 %) and SP (33.9 %), which raises concerns since mQTL are expected to be highly tissue-specific. We manually inspected the top 10 associations (i.e. with the highest absolute value of log_10_ odds ratio) and discovered additional unaccounted problems for 70 % (WB) and 40 % (SP), respectively (more details at **Supplementary Methods: Section III, Fig. S29**). Briefly, even though we removed CpGs affected by SNPs or indels, we still encountered G/H SNPs at CpG/CpH sites (H: A/C/T) and C/D SNPs at CpG/DpG sites (D: A/T/G), whose interpretation is even more challenging for the variant caller following bisulfite conversion. For example, for a T/C SNP at a CpG site, the C allele serves as a reporter of the methylation status of the region via co-methylation. However, if the region is variably methylated, the SNP caller may confuse the CpG epialleles for the SNP alleles post-conversion. This is especially problematic when the T allele is both the minor and the reference allele. Towards this end, 7 (11.3 %) and 12 (22.2 %) of the CpGs associated with genetic variants in WB and SP co-localized with JRCs^WB^ and JRCs^SP^, respectively.

### Systematic classification of jointly regulated CpGs with JRC_sorter

Given our limited success at verifying genetically driven changes in DNA methylation with pASM, we attempted a different strategy, inspired by our prior work looking into so-called epigenotypes (Planterose Jiménez et al. 2021); more specifically, clustering patterns in the unmethylated-methylated count plane (U/M-plane), where each point corresponds to a different individual (**Fig. 6A**). To name a few examples, imprinting is expected to generate a single intermediately methylated cluster (Model 1, M_1_) and so would cell type-specific regulation or even fully unmethylated or methylated regions across the cohort (which we do not expect since JRCs we pre-selected to be IMRs in the pool). Besides, polymorphic imprinting (i.e. imprinting at specific *loci* that is present in certain individuals but not others) is expected to generate two clusters, where the first corresponds to intermediately methylated, whilst the second may either be fully unmethylated (loss of methylation (LOM); model 3 (M_3_)) or methylated (gain of methylation (GOM); model 4 (M_4_)) (Monk et al. 2019). Alternatively, a strong co-dominant *cis*-mQTL caused by an underlying autosomal bi-allelic genetic variant is expected to generate three clusters: fully methylated, intermediately methylated and fully unmethylated, corresponding to homozygous to one allele, heterozygous and homozygous to the other allele (Model 5, M_5_). However, a *cis*-mQTL at some region in the Y-chromosome will solely generate fully methylated and unmethylated clusters on male individuals, since this region is effectively hemizygous (Model 2, M_2_). A more formal definition of the different models is available at **Supplementary Methods: Section V**.

**Figure 6.**
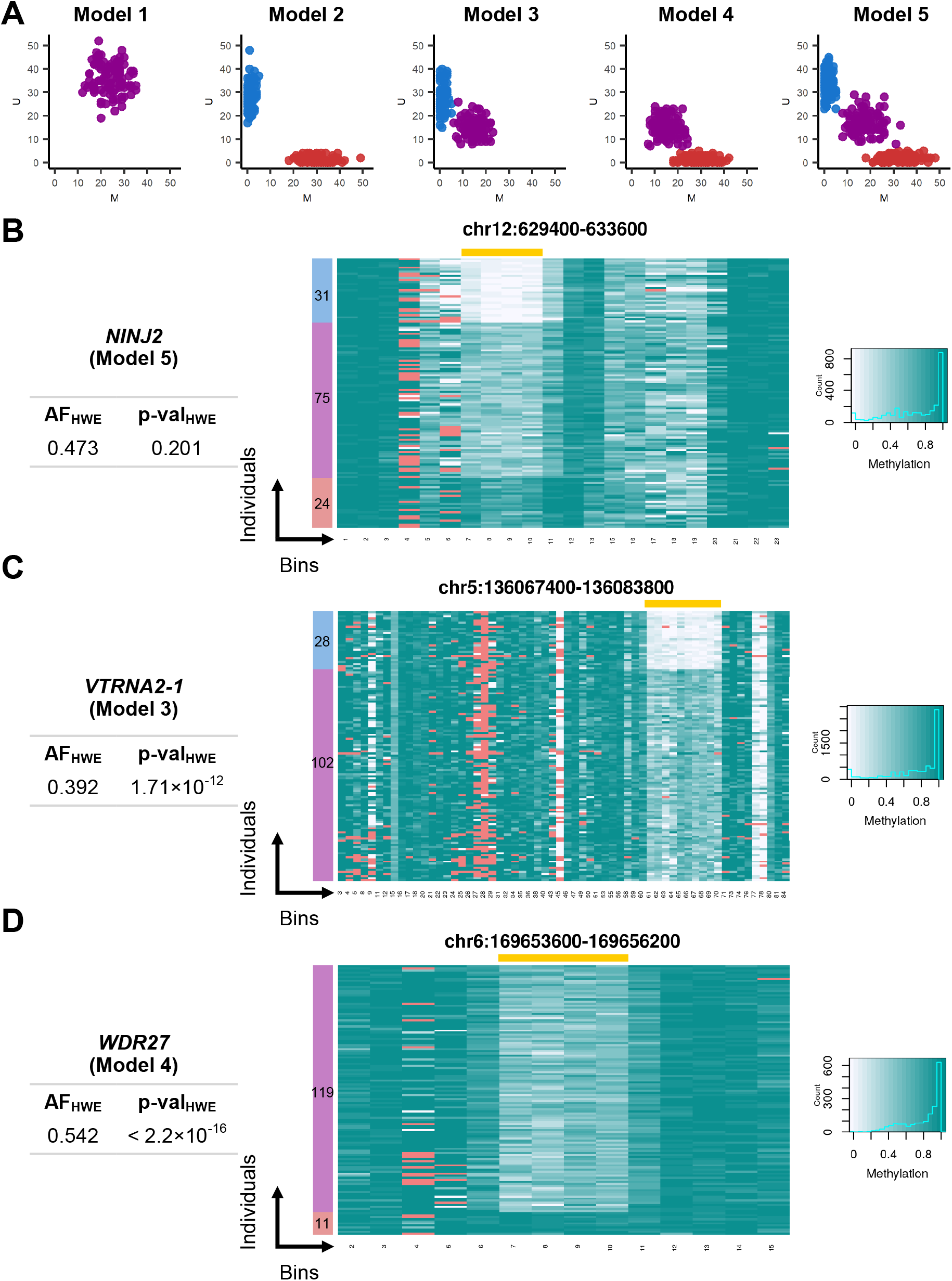
JRC classification with JRC_sorter. (A) The five epigenotype models considered by JRC_sorter. Heatmap of methylation levels at highlighted examples: (B) behaving as M_5_ from a verified mQTL at an intron in the *NINJ2* gene, (C) behaving as M_3_ at the polymorphically imprinted mono/bi-allelic switch at the *VTRNA2-1* gene and (D) behaving as M_4_ at an intron in the *WDR27* gene. Missing values are shown in red. Row colours (blue, violet and red) represent the different epigenotypes (unmethylated, intermediately methylated and methylated, respectively) at the active region of the JRC (highlighted in yellow). Examples (C) and (D) are inconsistent with HWE as shown on the summary statistics.

We hypothesised that assigning JRCs to epigenotype models could give us great insight into the underlying source of epigenetic variation. However, a pooled WGBS captures inter-haplotype, inter-cell and inter-individual variation but cannot distinguish between each other. Thus, to define epi-genotypes we require non-pooled WGBS data. Towards this end, we identified a publicly available non-pooled WGBS cohort dataset on cord blood from 130 newborns (62 typically developing and 68 later diagnosed with autism spectrum disorder (ASD)). which we purposed for the classification of JRCs^WB^ (**Fig. S30**). Importantly, we decided to include ASD samples in our analysis since this phenotype has minor, if not any, impact on the methylome (this is discussed further at **Supplementary Methods: Section IV**). Unfortunately, we did not find any suitable dataset to verify JRCs^SP^.

To automate the classification of JRCs, we developed JRC_sorter, a model-based classifier for JRCs (**Supplementary Methods: Section V**). JRC_sorter extracts methylation counts within a given JRC region across individuals and computes the likelihood under each model. We tested its performance based on a simulation study (**Fig. S31**, **Supplementary Methods: Section VI**). Briefly, we achieve almost perfect classification under our idealized simulation set-up given enough coverage (∼ 50 reads per CpG and sample), sample size (∼ 100 individuals) and allelic frequency (∼ 0.15). Moreover, since model M_5_ self-contains M_2_, M_3_ and M_4_, ties in model assignations are to be expected. Particularly, a *cis*-mQTL may in practice manifest as two clusters when the allelic frequency for its causal variant is low, since the sampling of the homozygous epigenotype for the minor allele becomes disfavoured at a finite sample size. For instance, we can distinguish this scenario from polymorphic imprinting since the latter should produce a missing epigenotype and excess heterozygosity. Thus, to break ties we tested for Hardy-Weinberg equilibrium (HWE) on the estimated epigenotype frequencies. From the 4215 JRC^WB^, JRC_sorter generated the following classification: 3286 (M_1_), 0 (M_2__M_5_), 70 (M_3__M_5_), 56 (M_4__M_5_), 45 (M_5_), while and 758 (inconclusive). We made available selected examples for M_1_ (**Fig. S32**), M_3_ or M_4_ (**Fig. S33**) and M_5_ (**Fig. S34**). We also note that X-inactivation in female samples drove certain JRCs in ChrX to be classified as M_3_ or M_4_ in the cohort (**Fig. S35**).

To highlight a few examples, the intronic mQTL at the *NINJ2* gene (Planterose Jiménez et al. 2021) (**Fig. 3A**) was confidently classified as M5 (**Fig. 6B**), as was the mQTL at the promoter of Peptidase M20 Domain Containing 1 (*PM20D1*), previously described for its role in Alzheimer’s disease (Wang et al. 2020) (**Fig. S34**). Regarding M_3_, we identified vault RNA 2-1 (*VTRNA2-1*) (**Fig. 6C**), a well-studied mono/bi-allelic switch that is polymorphically-imprinted: ∼75 % of individuals express the paternally-inherited allele (imprinted), while ∼25 % express *VTRNA2-1* bi-allelically (loss of imprinting) in a wide range of tissues (Silver et al. 2015; Carpenter et al. 2018). Its huge excess of intermediately methylated individuals is strongly inconsistent with a co-dominant biallelic *cis*-acting mQTL (p-val_HWE_ = 1.71×10^-12^). We verified these findings on DNA methylation microarray data (**Fig. S36**). Other examples of M_3_ behaviour include regions at the promoters of Dual Specificity Phosphatase 22 (*DUSP22)*, Cytochrome P450 2E1 (*CYP2E1)* and Olfactory Receptor Family 2 Subfamily L Member 13 (*OR2L13)*. Together with *VTRNA2-1*, these four M_3_-behaving genes were highlighted as putative metastable epialleles in a prior study (Van Baak et al. 2018).

Moreover, an intronic region at the tryptophan-aspartic acid (WD) Repeat Domain 27 (*WDR27*) gene (**Fig. 6D**) displayed clear M_4_ behaviour (p-val_HWE_ < 2.2×10^-16^). Notably, this region was also a JRC^SP^ (**Fig. S37**). Looking into DNA methylation microarray data on both WB and SP, we confirmed that this region behaves as M_4_ in WB, but intriguingly discovered that it behaves as M_5_ in SP. This puzzling observation can be explained by a simple model: this JRC is an mQTL in SP. Diploid spermatogonia display three possible epigenotypes (fully methylated/unmethylated or intermediately methylated) depending on whether a bi-allelic genetic variant is homozygous for one allele, for the other or heterozygous, respectively. However, upon meiosis, haploid sperm cells are generated: for example, heterozygous spermatogonia generate approximately 50 % of sperm carrying the M or the U epiallele; the characteristic trimodal methylation distribution of M_5_ still arises from haploid population of cells since we assay DNA methylation in bulk. Contrarily, we hypothesise that egg cells persistently carry the M epiallele (i.e. the mQTL is not functional in oocytes). Upon fertilization, resulting zygotes will present a fully methylated or intermediately methylated epigenotypes at this *locus* (i.e. M_4_), depending on whether the fertilizing sperm cell carried the M or the U epiallele. Exploring the implications of this model, methylation epigenotypes in the zygote must then survive epigenetic reprogramming during early development to be able to observe a bi-modal methylation distribution in WB. If true, we should observe bimodal methylation distributions for other tissues; we indeed confirmed this prediction for a wide range of tissues but not in placenta (**Fig. S38A**). Moreover, there must be paternal imprinting; this has been previously shown in the literature (Monk et al. 2019). Additionally, we estimated the allelic frequency of the underlying causal variant taking into account this model, obtaining 0.085 (cord blood, USA, WGBS, n = 140) and 0.113 (SP, USA, microarray, n = 379), which were not significantly different (p-value = 0.446; Fisher’s exact test). We also estimated allelic frequencies in other populations: 0.154 (WB, Sweden, microarray, n = 727) and 0.104 (WB, the Gambia, microarray, n = 240); differences of which could be explained by sampling error (Fisher’s exact test, p-value = 0.055) (**Fig. S38B**). Lastly, we found evidence of *cis*-acting genetic variants in WB from mQTLdb (Gaunt et al. 2016) and proposed candidate *cis*-acting causal genetic variants regulating this *locus* (**Fig. S39**).

## Discussion

In the post-GWAS era, epigenomics picks up the baton in the phenome-mapping relay race. The genetic component of epigenetic inter-individual variation may shed light on the underlying mechanisms of action for discovered genetic biomarkers, whilst the environmental component may serve as a proxy to study the non-heritable part of human phenotype. However, the human genome makes up a gargantuan search space, so novel approaches to affordably highlight regions of interest from its vast genomic territory are in high demand. Yet, traditional approaches to map methylome inter-individual variation have limitations of their own. EWAS requires matching reliable and identifiable phenotypic information and struggles with confounders. mQTL mapping requires matching genotypic data and suffers from huge redundancy due to LD (linkage disequilibrium) between genetic variants and co-methylation of neighbouring CpGs. Both are traditionally carried out on DNA methylation microarrays that cover solely 1.5-3 % of the methylome with associated challenges such as background fluorescence, batch effects, genetic artifacts or cross-hybridizing probes. Alternatively, ASM/pASM is typically performed on WGBS data, but can only discover imprinting sites/mQTL at read-length resolution. More critically, it requires variant calling of heterozygous genetic variants (when matching genetic data is unavailable), which is highly unreliable at low-coverage WGBS data. For example, the putative *cis*-causal variant for the mQTL at the *NINJ2* gene coincides with a substantial read coverage drop (**Fig. S13**). Reliably distinguishing variants from sequencing errors with only a handful of bisulfite reads is still a challenging computational problem.

Under this setting, we developed Binokulars, a novel computational approach for identifying regions under active methylation regulation given pooled WGBS data, which operates without requiring matching phenotypic/genotypic data. It does so by detecting the signatures of CWR via a genome-wide randomization test. Its success ultimately relies on the natural processivity of the enzymes that regulate the methylome (DNMT1, DNMT3A, DNMT3B and TET). Similar concepts to CWR have been previously proposed (Shoemaker et al. 2010; Landan et al. 2012; Guo et al. 2017; Jenkinson et al. 2017; Orjuela et al. 2020; Scott et al. 2020; Ding et al. 2022); in such studies, authors typically compute some sort of summary statistic (e.g. entropy, methylation LD, epi-polymorphism) and use empirical thresholds to highlight certain regions. In other words, unlike Binokulars, they do not control for false discovery (whether observations could have arisen by chance). Also, none of these previous studies have been carried out using pooled data, where this mapping approach meets its full potential. Towards this point, Binokulars’ effectiveness lies in its generality: this approach can detect the signatures of CWR in regions affected by (polymorphic) imprinting, cell type and tissue-specific regulation, *cis*-and *trans*-mQTL or associated with (un)known phenotypes as long as variation of which is present in the pool. In fact, it would be possible to combine healthy and diseased individuals in the pool to also identify JRCs associated with disease-specific deregulation. Strikingly, all findings in this study derived from pooled data on solely 12 individuals. This observation reveals how Binokulars makes the most value out of the data by working at the deepest level (e.g. reads). This let us wonder about the number of new findings for both health-related and disease-associated traits that may arise from future studies implementing this strategy on larger pools across tissues.

At the same time, in its generality lies its limitation. The pooled WGBS experimental design confounds inter-haplotype, inter-cell and inter-individual methylome variation. Binokulars successfully identifies JRCs but cannot reveal the underlying factors that mediates active regulation. Because of this, we additionally developed JRC_sorter, a novel computational approach for classifying JRCs into five different epigenotype models given population-level WGBS data. With this approach and cohort WGBS data on cord blood, we were able to shed light onto JRCs^WB^ associated with mQTL and polymorphic imprinting. Specifically, we highlighted two examples of polymorphic imprinting: *VTRNA2-1* (M3) and *WDR27* (M4); loss of imprinting for these sites results from environmental (i.e. availability of methyl donors during early development (Silver et al. 2015)) and genetic causes (*cis*-mQTL in SP), respectively. As for SP, no suitable dataset could be identified at the time to classify JRCs^SP^ with JRC_sorter, limiting our exploration. For example, the JRC^SP^ at the *PLEC* gene could not be attributed to any known source of epigenetic variation. In general, in spite of being a human haploid model with huge implications in *trans*-generational epigenetic inheritance and imprinting, this tissue continues to be understudied. For instance, to the best of our knowledge, no mQTL mapping study has been carried out on SP yet.

Next, we validated our approach by comparing our outcomes with traditional methodologies. On the one hand, our comparison with pEWAS was fruitful and helped attribute JRCs to known sources of epigenetic variation such as tissue-specific regulation or ageing. On the other hand, we struggled to extract meaningful information from pASM due to genetic artifacts caused by SNPs and indels, in contrast to the original study from Kaplow *et al* that pioneered this strategy (Kaplow et al. 2015). To explain the differences, Kaplow *et al* performed a total of 823,726 comparisons as supposed to our case, where we tested ∼53 M. Looking closely into their work, this drastic reduction probably results from the pre-masking of the reference genome at positions with known SNPs (MAF > 0.04) prior to alignment, which effectively removed CpG-SNPs from their data. Additionally, Kaplow *et al* set their genome-wide significance threshold at α=0.001, obtaining 2379 significant hits. We were considerably more stringent by applying Bonferroni multiple testing correction. Thus, the aforementioned methodological differences explain the different outcomes of our studies. In general, a common issue across all methodologies implemented here (pASM, pEWAS, Binokulars) are false discoveries resulting from low-mappability regions. For example, even after filtering out regions with at least one random mappable 100-mer as defined by BisMap (Karimzadeh et al. 2018) and removing reads with mapping quality < 40 (default behaviour of BISCUIT), dubious highly significant JRCs were still identified in close proximity to low-mappability regions (**Fig. S41**). Further improvements towards this end are still required.

Profoundly, our findings inspired the following question: how do processive enzymes give rise to non-jointly regulated CpGs at intermediately methylated regions? At low CpG density regions, when the mean processive length is substantially smaller than the inter-CpG distance, little information is carried on from CpG to CpG, resulting in quasi-independent regulation. To what extent CpG-poor regions contribute to the functional regulation landscape is hard to say. For instance, these regions are typically of later replication timing and of poorer DNA methylation maintenance (Zhou et al. 2018; Petryk et al. 2021; Higham et al. 2022); thus, relying on such regions for proper genomic regulation may be less robust. On the contrary, the emergence of non-jointly regulated CpGs at intermediately methylated regions at CpG-rich regions is harder to envision. We speculate that one way to induce methylation epihaplotype decoherence could be by simultaneous recruitment of DNMT and TET; under a scenario of constant writing/re-writing, CWR decreases with every new cellular generation. For example, CWR is typically not present at sharp junctions between unmethylated and methylated domains (**Fig. S41**). Such junctions are maintained by recruitment of DNMT1/TET at either side of the boundary. Thus, simultaneous recruitment of DNMT and TET is expected where these two domains meet, thus potentially favouring decoherence as supposed to the formation of a JRC.

Lastly, highlighting regions of interest in the methylome can be helpful in the design of future DNA methylation microarrays. Having this in mind, we evaluated the representation of JRCs in two microarray platforms (Infinium HumanMethylation450 and MethylationEPIC beadchip) by counting the number of probes that co-localized with JRCs^WB^ and JRCs^SP^ (**Fig. S42**). Our comparison revealed a better probe coverage in WB compared to SP. In general, the effects of microarray design bias remain unexplored and are particularly challenging to unveil since manufacturers tend to seclude from transparency at product development stages for intellectual property safety reasons. Yet, their choices dictate the initial pool of markers in EWAS and thus, determine whether this approach successfully identifies markers of interest or not, unavoidably defining the whole methylomics field.

## Methods

### Theory of Binokulars and simulations

Binokulars is a randomization test for CWR from pooled WGBS data. It tests whether a given region displays a tendency for CpGs to display the same epiallele (unmethylated/methylated) at a higher level than what it is expected from a binomial distribution (i.e. whether they are JRCs) (**Fig. 1A-B**). A formal account on the theory of the method is available at **Supplementary Methods: Section I**. Briefly, Binokulars bins an input region, estimates the methylation ratio per bin, counts the number of methylated and unmethylated cytosines at CpG sites per read and bin to finally compute a test statistic: the negative log-likelihood (“surprise”) of the data under a binomial model. Subsequently, Binokulars samples the null distribution of the test statistic via bootstrapping to obtain a randomization p-value. Given the large computational cost per permutation and in order to withstand a genome-wide multiple testing burden, Binokulars also generates approximate p-values based on a parametric assumption whenever the randomization p-value is equal to zero. To compute this, we assume that the test statistic under the null distribution follows a Gamma distribution (though others propose alternatives in a different context (Knijnenburg et al. 2009)), the parameters of which are estimated from bootstrapped values for the test statistic.

To assess the limitations of Binokulars, we performed simulations with the following set-up (**Supplementary Methods: Section II**). Briefly, we defined a region of length *L* ∈ ℕ with CpG density of *ρ*_CpG_ ∈ ℝ^+^ and methylation ratio *p* ∈ [0,1], for which a total of *n*_reads_ ∈ ℕ are available of length *l* ∈ ℕ. We defined the effect size of the region *φ* ∈ [0,1] as the ratio of jointly regulated reads (reads where all cytosines bear the same epiallele) with respect to the total. A simulation begins by randomly placing CpGs in the region, respecting its dinucleotide nature (there cannot be a CpG at site *i* and *i* + 1). It then places all *n*_reads_ reads within the region with uniform probability. For all reads that overlap with CpG sites, it decides whether the read is “jointly regulated” or not via Bernoulli trial with probability *φ*. If the read turns out to be “jointly regulated”, a single Bernoulli trial with probability *p* is performed and the outcome of which (0 or 1) is set as the methylation status (U or M) for all CpGs contained within the read. If the read does not turn out to be “jointly regulated”, independent Bernoulli trials with probability *p* are performed for each CpG contained within the read. The following parameters were fixed: *L* = 2000 bp, for convenience; *l* = 100 bp, to fit the typical WGBS read length. Binokulars was executed with a bin size *δ* = 200 bp, to fit nucleosomal resolution, number bootstrapping iterations *N*_*boot*_ = 1000 and flank length, *f* = 0; we instead tested ranges of values for parameters *p*, *ρ*_CpG_, *n*_reads_ and *φ*. To select a meaningful range for *ρ*_CpG_, we first computed CpG density across the entire human reference genome (hg38) (**Fig. S1**).

### Theory and data for discovery of jointly regulated CpGs with JRC_seeker

JRC_seeker is a Snakemake pipeline (Köster and Rahmann 2012) for genome-wide discovery of JRCs (**Fig. S2**). We developed it on WGBS data (directional library) from pooled (equimolar) publicly available WB and SP samples derived from healthy males (Laurentino et al. 2020). In total, four pairs of paired-end fastq files were made available by Laurentino *et al*: ERR2722068 (WB; pool of 6 individuals; aged 61-71; 15.66x), ERR2722069 (SP; pool of 6 individuals; aged 61-71; 16.32x), ERR2722070 (WB; pool of 6 individuals; aged 18-24; 19.29x) and ERR2722071 (SP; pool of 6 individuals; aged 18-24; 16.34x). We concatenated fastq files *in silico* with *cat* to generate three pairs of fastq files (i.e. paired-end): WB (by combining ERR2722068 and ERR2722070; pool of 12 individuals), SP (by combining ERR2722069 and ERR2722071; pool of 12 individuals) and WB+SP (by combining all fastq files). The main input for JRC_seeker is an alignment file (.bam). We generated ours with BISulfite-seq CUI Toolkit (BISCUIT) (Zhou 2022), by marking duplicate reads with *samblaster* and aligning to the human reference genome (*hg38/GRCh38*) with *biscuit align* in paired-end mode.

To begin the pipeline, JRC_seeker generates a pile-up (VCF file) with *biscuit pileup*, calls DNA methylation at CpG sites with *biscuit vcf2bed* with argument -t “cg”, and merges forward and reverse cytosine with *biscuit mergecg*. It then uses custom python scripts to generate binned binary methylated and unmethylated tracks ready for input to ChromHMM’s *learnModel* (Ernst and Kellis 2017). We employed a bin size of 200 bp (roughly nucleosomal resolution) and considered bins as intermediately methylated when methylation ratio lied between 0.2 and 0.8. For our multivariate HMM parameter estimation, we set the number of iterations to 500 and number of states to four, which generally correspond to fully methylated/unmethylated (10 or 01), intermediately methylated (11) and no data (00); though, this relationship is not strictly imposed. Since our hidden states encode for the same information as our observable states, HMM acts as a noise reduction algorithm rather than an integrator of combinatorial and spatial patterns. In any case, ChromHMM estimates transition and emission matrices and generates a maximum *a posteriori* probability estimate of the most likely sequence of hidden states, which we here denote as a genome segmentation. JRC_seeker then extracts IMRs from this genomic representation. Since some IMRs tend to be small, disjoint and sparse, we polished ChromHMM’s representation with a custom R-script, *binPolish*: based on *GenomicRanges*, we extended IMRs with flanks on both sides, removed low-mappability regions (as defined by Bismap (Karimzadeh et al. 2018)) and regions known to display anomalous coverage (blacklisted regions, as defined by ENCODE; annotations ENCSR636HFF and ENCSR797MUY), merged overlapping IMRs and removed any regions below 200 bp.

Having an IMR list, JRC_seeker can subsequently start processing reads. To do so, it parses the bam file to the epiread format with *biscuit epiread*, which previously requires calling SNPs with *biscuit vcf2bed* argument -t snp. We then used a custom bash script to further combine, compress and split per chromosome for increased querying performance. The resulting formatting is a tab-separated file where each line corresponds to a read that contains CpG sites, with the following fields: chromosome, start position, number of methylated and unmethylated cytosines in forward and in reverse position. For each IMR, JRC_seeker calls *awk* to extract the reads from this file, whose start position lies within the IMR region with default flanks *f* = 500 bp and executes Binokulars, with default bin size *δ* = 200 bp and number bootraps *N*_*boot*_ = 1000.

### Functional annotation of jointly regulated CpGs

A list of imprinted genes was obtained from Geneimprint (Jirtle 2012). Summary EWAS statistics and raw microarray data on WB cell-type composition were accessed via the FlowSorted.Blood.450k R-package and commands *data(FlowSorted.Blood.450k. compTable)* and *data(FlowSorted.Blood.450k)* (Reinius et al. 2012). Manhattan and QQ plots were obtained with *qqman::manhattan* and *qqman::qq*, respectively. To assign genic features to JRCs, we employed the gene annotation file “hg38.refGene.txt” obtained from UCSC. We defined “promoter” as the region spanned 1 kb from transcription starting site. In case of category conflict, the following priority relations were established: promoter > exon > intron > intergenic. Enrichment tests were performed with *stats::fisher.test*. Gene list functional enrichment was performed with *gprofiler2::gost* on ontologies gene ontology (GO), Kyoto encyclopaedia of genes and genomes (KEGG), reactome (REAC), transfac (TF), micro-RNA (MIRNA), human protein atlas (HPA), protein complexes (CORUM), human phenotype (HP) and WikiPathways (WP).

### Pooled epigenome-wide association study

Firstly, using *samtools view* and *grep*, we split the bam files that were previously employed in JRC discovery into their corresponding components: WB (ERR2722068, ERR2722070) and SP (ERR2722068, ERR2722070). We performed methylation calling on each sub-bam files like within JRC_seeker: *biscuit pileup*, *biscuit vcf2bed -t cg* and *biscuit merge cg*. The obtained bed files were input and parsed into the R-programming environment. Specifically, we used functions from the DSS R-package to perform EWAS in absence of technical replicates by harnessing genomic spatial information. To do so, we called *DSS::DMLtest* with parameters smoothing=TRUE, equal.disp=FALSE and smoothing.span = 500; and *DSS::callDMR* with parameters delta=0.1, p.threshold=1e-5, minlen=200, minCG=5, dis.merge=200 and pct.sig=0.5. Differentially methylated regions (DMRs) were visualized with *DSS::showOneDMR*. A total of three pEWAS were performed: WB (old and young) *versus* SP (old and young), WB (old) *versus* WB (young) and SP (old) *versus* SP (young). We also defined the pEWAS background sets by calling *DSS::callDMR* with parameters delta=0, p.threshold=1, minlen=200, minCG=5, dis.merge=200 and pct.sig=0.5. For the colocalization analysis, we additionally trimmed DMRs from low-mappability and anomalous-coverage regions, as performed above. For colocalization analysis, we visualized the interactions between the different sets as a network using the GGally, ggnet, network and sna R-packages. We established the node side to represent the number of megabases spanned by any given genomic set and the edges to represent the overlapping megabases between the different pairs of genomic sets.

### Pooled allele-specific methylation

We performed pASM in WB (ERR2722068+ERR2722070; pool of 12 individuals) and SP (ERR2722069+ERR2722071; pool of 12 individuals) with BISCUIT (Zhou 2022). Corresponding alignment files (bam files) were converted into pile-ups (vcf files) with *biscuit pileup*. Variant calling was performed from the pile-up with *biscuit vcf2bed -t snp*. To run ASM, we firstly parsed the pile-up into the pairwise epiread format with *biscuit epiread -p* and then ran *biscuit asm*, which computes Fischer’s exact test for all pairs of variably methylated CpGs/heterozygous SNPs that were jointly profiled at the read-level. However, a large percentage of associations are compromised for several reasons. We excluded associations for called variants that were not registered in dbSNP155 with a minor allele frequency (MAF) of at least 0.05; SNP positions and allele frequencies were obtained from the dbSnp155Common.bed file at UCSC. We removed genetic artifacts by excluding associations in which the CpG co-localizes with SNPs/indels (MAF > 0.05); we extracted CpG positions from the BSgenome.Hapiens.UCSC.hg38 R-package with *Biostrings::matchPattern*. Moreover, we removed associations where the CpG lied within low-mappability and anomalous-coverage regions, as performed above. Following exclusion of the aforementioned associations, we selected at Bonferroni significance threshold (1.28×10^-8^ (WB) and 1.69×10^-8^ (SP)) and for DNA methylation differences of more than 0.2 for one allele over the other.

### Theory and data for jointly regulated CpG classification with JRC_sorter

A formal exposition on the theory of the JRC_sorter is available at **Supplementary Methods: Section IV**. Briefly, unlike Binokulars, JRC_sorter does not employ read level information but instead the unmethylated/methylated count matrices (per CpG andindividual). We developed it for the classification of JRCs^WB^ with non-pooled WGBS data on cord blood from 62 typically developing newborns and 68 newborns diagnosed with autism spectrum disorder (ASD) at 36 months. Instead of processing reads from scratch, we directly used the publicly available processed methylation reports provided by Mordaunt *et al* (Mordaunt et al. 2020). These were generated by aligning over 2 Tb of read data to the hg38/GRCh38 reference genome with Bismark. To proceed, we first parsed all 130 methylation reports using *awk*, *sed* and *bedtools intersect* to obtain a tab-separated file, split per chromosome, where each line corresponds to a CpG, with the following fields: chromosome, start and end position, number of methylated and unmethylated cytosines in forward and in reverse position.

Briefly, per JRC, JRC_sorter first constructs the unmethylated and methylated count matrices across individuals for CpGs within the JRC by extracting the cytosine counts from each parsed file with *awk*. It then aggregates the count matrices into bins (bin size, *δ* = 200 bp) and computes likelihoods per bin for five distinct models (M_1_-M_5_). We offset fully methylated and unmethylated states by *ε* = 0.05 to permit measurement error. JRC_sorter assigns each bin to the model with the maximum log-likelihood, but allowing certain tolerance (i.e. bin tolerance, ψ = 0.15) to enable ties (e.g. bin assignation). At this stage, any ties that include the M_1_ model are simply resolved as M_1_ (e.g. the simplest model is favoured). To compute JRC-level likelihood for model Mi, we sum the log-likelihoods across bins: for bins that include label Mi, JRC_sorter uses M_i_ log-likelihood, while for all other bins, it uses the M_1_ log-likelihood. This way, JRC_sorter is tolerant to misspecification of the precise JRC coordinates. Extending this procedure to all JRCs, JRC_sorter finally generates the log-likelihoods for all five models across JRCs. To classify JRCs into different models, we select the model that produces the maximum log-likelihood but again, allowing certain tolerance (i.e. model tolerance, *κ* = 0.1) to enable ties (e.g. model assignation). Since model M_5_ encapsulates models M_2_, M_3_, M_4_, ties such as M_2__M_5_, M_3__M_5_, M_4__M_5_ are to be expected by definition. To resolve them, we employ a statistical test for Hardy-Weinberg equilibrium.

### Validation using DNA methylation microarray data

WB and SP DNA methylation microarray data from public GEO resources were used for validation (Johansson et al. 2013; Van Baak et al. 2018; Pilsner et al. 2022), whilst data on isolated cell types was accessed via the FlowSorted.Blood.450k R-package (Reinius et al. 2012). To pinpoint genetic variants to the *WDR27* polymorphic imprinting site, we used WB mQTL summary statistics from mQTLdb (Gaunt et al. 2016). To access DNA methylation distributions across a large panel of tissues, we used the API platform from EWAS datahub (Xiong et al. 2020).

## Data Access

WB and SP WGBS data is available at ENA under accession number PRJEB28044 (ERR2722068, ERR2722069, ERR2722070, ERR2722071) (Laurentino et al. 2020). Cord blood WGBS data are available at GEO under accession number GSE140730 (Mordaunt et al. 2020). WB and SP microarray data is available at GEO under accession numbers GSE185445 (Pilsner et al. 2022) (SP) and GSE87571 (Johansson et al. 2013); GSE99863 (Van Baak et al. 2018) (WB). Isolated WB cell type microarray data is available via the FlowSorted.Blood.450k R-package (Reinius et al. 2012). To access DNA methylation distributions across large panel of tissues we used the API from EWAS datahub (Xiong et al. 2020). As for the our developed software we have been made available four Github repositories: data analysis R-scripts (https://github.com/BenjaminPlanterose/JRC_downstream_analysis, Binokulars (https://github.com/BenjaminPlanterose/Binokulars), JRC_seeker (https://github.com/BenjaminPlanterose/JRC_seeker) and JRC_sorter (https://github.com/BenjaminPlanterose/JRC_sorter).

## Acknowledgements

We thank all researchers that made their data publicly available, without whom our study would not have been possible. The work of all authors is funded by Erasmus MC. AV is additionally supported by an Erasmus MC fellowship 2020.

